# Melchior: A Hybrid Mamba-Transformer RNA Basecaller

**DOI:** 10.1101/2025.01.11.632456

**Authors:** Elon Litman

## Abstract

Sequence transduction from raw nanopore signals is notoriously difficult because the signal level does not naturally correspond to a single base, but rather many adjacent nucleotides. Thus, we introduce Melchior, an RNA basecaller that uses a hybrid Mamba-Transformer backbone to achieve global visual context at a lower computational complexity. This is in contrast to temporal convolutions, which collate features by fusing both spatial and channel features in the local receptive field. Melchior is also able to exploit the full complementarity between local and global features, unlike Vision Transformers, which have empirically been observed to ignore local features. Augmenting a selective structured state-space sequence model with self-attention unlocks unprecedented performance gains, particularly in homopolymer regions, by modeling fine-grained details in both short and long-range spatial dependencies.

## 1 Introduction

Third-generation sequencers promise to revolutionize RNA and DNA sequencing through their generation of very long single-molecule reads, tens or hundreds of times longer than next-generation solutions. Oxford Nanopore Technologies (ONT), with the introduction of MinION devices, have realized reads with lengths up to 2.4 Mbp using nanopore sequencing. The nanopore sequencing paradigm is based on measuring ionic current in a biological membrane when a polynucleotide strand passes through an embedded channel. The ion currency values are then translated to a sequence of nucleobases through the process of basecalling.

However, sequence transduction from raw nanopore signals is complicated by several factors. Firstly, the signal level does not naturally correspond to a single base, but rather several nucleotides that are in the nanopore simultaneously. The translocation of homopolymers also does not change the underlying signal, so only signal changes can be used to disambiguate adjacent bases. Lastly, the non-uniform speed of the nucleobases passing through the nanopore makes the number of signal changes an inaccurate estimate of sequence length.

### 1.1 Related Work

Although ONT has officially created several basecallers, the specifics of their models have not been disclosed publicly. Consequently, various third-party basecallers utilizing deep learning have emerged, each employing different methodologies. Basecallers evolved from statistical tests, to hidden Markov models (HMMs), and finally to the use of neural networks. Mincall used a deep convolutional neural network (CNN) with residual connections to extract features from the raw signal [1], and URNano used a convolutional U-net with integrated RNNs [2]. Bonito is a deep learning-based basecaller recently developed by ONT [3]; its architecture is composed of a single convolutional layer followed by three stacked bidirectional gated recurrent unit (GRU) layers. Although Bonito has achieved relatively high accuracy, its speed is too slow to be used in production. Most recently, Halcyon used a sequence-to-sequence (Seq2Seq) model with attention [4]. In practice, attention and RNNs are applied in conjunction with CNNs or used to replace certain components of CNNs while keeping their overall structure in place [5]. In order to handle the variable-length output dimension, a connectionist temporal classification (CTC) decoder is usually used [6]. This technique, which has historically been used to process speech signals [7], was prominently featured in the third-party basecaller Chiron [8].

However, basecallers built solely with attention have not yet demonstrated higher accuracy for basecalling [4]. This is thought to be because the nanopore signals have more vague transitions between nucleotides than words in natural language processing tasks, especially within homopolymers or repeats consisting of purines or pyrimidines [9]. We posit that current architectural paradigms face fundamental limitations in processing nanopore signals: convolutional networks are inherently constrained by their local receptive fields, precluding access to global context, while attention-based architectures, despite their theoretical ability to capture long-range dependencies, exhibit both computational inefficiency and empirically poor synthesis of local features. These limitations are particularly salient in basecalling, where models must simultaneously reason about fine-grained signal characteristics and broader sequence context. Current approaches, constructed on these primitives, operate independently of prior biological and physical insights, leading to suboptimal feature extraction that fails to fully leverage the inherent structure of nanopore signals.

### 1.2 Theoretical Considerations

A critical point in the development of new architectures has always been the receptive field, that is, the area of the input on which the output values depend. The two most popular backbone networks in visual representation learning are deep CNNs and Vision Transformers (ViTs). Although CNNs can effectively extract local features, they have difficulty capturing global context and long-range dependencies. Various methods, such as dilated [10] or deformable [11] convolutions, attempt to enlarge the receptive field while maintaining the original inductive biases and translation invariance. In practice, however, the field remains mostly limited to (semi-)local areas. Squeeze-and-excitation (SE) blocks [12] sought to enhance the representational power of deep CNNs by learning to selectively emphasize channel-wise feature responses and suppress less useful ones. Notably, RODAN utilized squeeze-and-excitation in an end-to-end CNN to facilitate global processing in the basecalling task [13]. Among the limitations of the SE block, arguably, is the squeeze operation that performs global information embedding. The global average-pooling used to compute channel-wise attention is deleterious to local features, which may be necessary for identifying the importance of different channels [14]. Indeed, the SE module also misses the entire motivation of spatial attention, referring to the ability of the network in deciding “where” to focus.

[15] introduced the ViT, adapting the concept of self-attention to achieve a global receptive field by processing non-overlapping image patches. Compared to CNNs, ViTs generally demonstrate superior learning capabilities on large-scale data; however, the quadratic complexity of self-attention w.r.t. to the number of tokens introduces substantial computational overhead in downstream tasks involving large spatial resolutions. This can be prohibitive in basecalling, where a signal may contain thousands of time steps. Swin Transformer improves efficiency by adopting non-overlapping windows and slowly increasing the receptive field by window shifting to describe interactions between different stages. As a result, the ability of the self-attention to capture long-range information is limited. Using the original columnar structure of the ViT facilitates the processing of long-range dependencies, but also deteriorates for local feature details [17]. Approaches that enable a ViT to attend to fine-grained global and local features have been described, usually with added architectural complexity and computational costs [18, 19]. Nevertheless, an efficient treatment to fully exploit the complementarities of global and local representations in the nanopore signal remains to be elaborated. Thus, a novel architecture may be prophylactic for the basecalling task.

Mamba [20] proposed a new State Space Model (SSM) that achieves linear time complexity and excellent performance in modeling long-range dependencies. The core contribution of Mamba is the merging of time-varying parameters into SSMs and the development of a hardware-aware selection mechanism for efficient training and inference. This architectural innovation was used as the basis of Caduceus, a family of bidirectional long-range DNA sequence models that excel at a wide range of predictive tasks in genomics [21]. Despite Mamba’s strong theoretical guarantees and advantageous deployment characteristics, its autoregressive formulation is inimical to vision applications. VMamba [22] pioneered a vision backbone that relies on a cross-scan module to reconcile the ordered nature of 1D selective scan and the non-sequential structure of vision data. Similarly, Vim [23] incorporated position embeddings into a bidirectional SSM for global visual context modeling. Such bidirectional SSMs introduce significant latency, as they require processing the entire sequence before making predictions. This added complexity also entails training challenges, increases the risk of overfitting, and does not consistently improve accuracy. Therefore, Transformers and deep CNNs remain preferred for basecalling.

Recently, Vision Mamba described MambaVision Mixer [24], a simpler block that can capture both short and long-range visual dependencies. They also studied different isoparameter integration patterns to find that self-attention in the neck significantly enhances the network’s capabilities. The resulting hybrid architecture comprising MambaVision Mixer and Transformer blocks achieves a new state-of-the-art Pareto front on the ImageNet-1K dataset in terms of Top-1 accuracy and image throughput tradeoff. We consider that nanopore data could especially benefit from a hybrid Mamba-Transformer architecture, which could be constructed to improve over monolithic approaches while maintaining the original efficiency of SSMs. Such a model could account for nucleobases acting at a distance on the signal corresponding to a given base through location-aware global context understanding, while gleaning granular features relating to homopolymer translocation and the unique structure of RNA. We focus on RNA rather than DNA basecalling, for the former being rarely studied in comparison.

### 2 Methodology

### 2.1 Problem

Formally, an input with a *T*-timestep signal is denoted by ***s*** = [*s*_1_ … *s*_*T*_]. We seek an *N*-nucleobase sequence, represented by ***Y*** = [***y***_1_ … ***y***_*N*_] where ***y***_*k*_ ∈ ℝ^5^, for the probabilities of the four canonical nucleotides and the IUPAC ambiguous base (A, T, G, C, and N) at position 1 ≤ *k* ≤ *N*.

### 2.2 Macro Architecture

As illustrated in Figure 1, Melchior has an isotropic structure consisting of three stages: the stem, backbone, and head of the network. The stem allows for fast feature extraction by applying a series of 1D convolutions followed by batch normalization and Gaussian Error Linear Unit (GELU) activations to transform the input signal. The backbone is comprised of *N* = 20 alternating MambaVision Mixer and Transformer blocks with residual connections. Self-attention recovers lost global context and improves the capture of long-range dependencies. Lastly, the head truncates the sequence length by applying a 1D adaptive average pooling operation before projecting to the class scores through a fully-connected linear layer.

**Figure 1:**
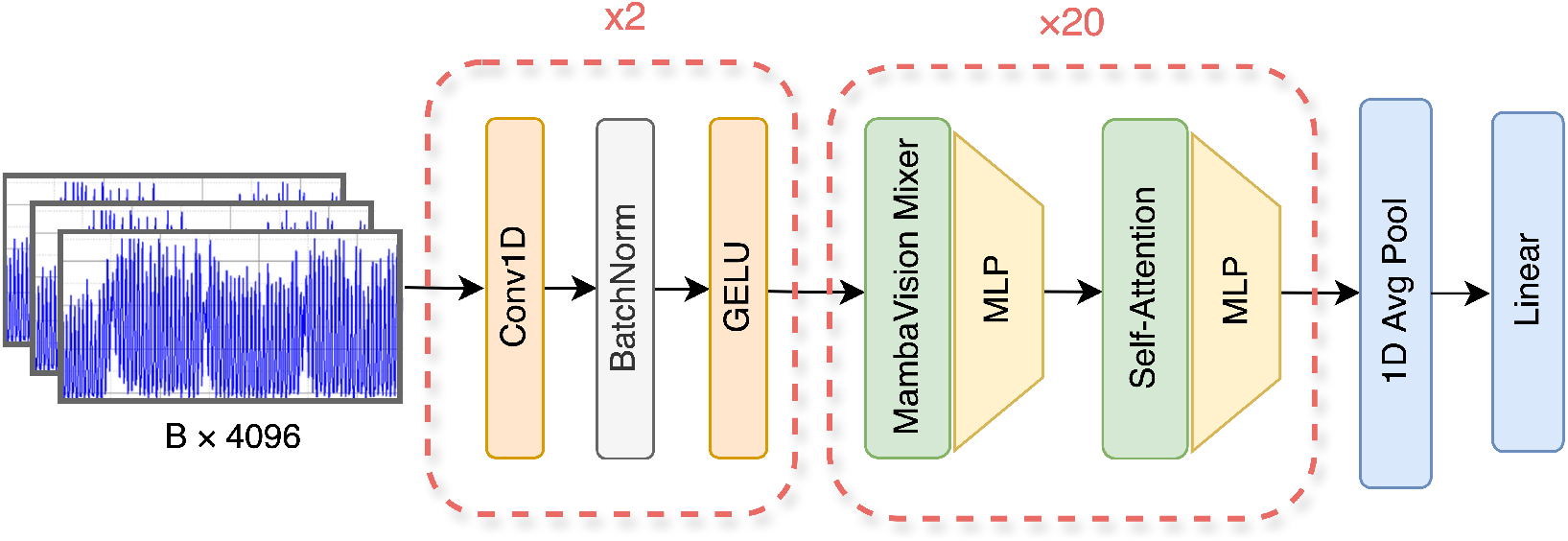
The architecture of Melchior is comprised of a fast feature extraction network at the stem, a hybrid Mamba-Transformer backbone, and head that projects to the output scores.

#### Stem

The stem operates on single-channel input signals with 4096 time steps. Rather than patchify the raw nanopore signal, we opt to embed the inputs using a fast feature extraction network. This subnetwork consists of two sequential 1D convolutional layers, each followed by batch normalization and GELU activation:

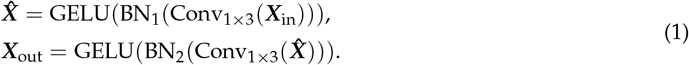

Here, Conv_1_×_3_ denotes 1D convolution operations with kernel size 3 and a padding of 1, BN_*i*_ represents the *i*-th Batch Normalization layer, and GELU is the Gaussian Error Linear Unit [25] activation function. Batch normalization [26] stabilizes the training process by normalizing the activations. Given an input batch ***X*** = [***x***_1_, …, ***x***_*n*_], where *n* is the minibatch size, the batch normalization layer calculates the mean value within the minibatch as

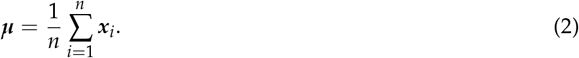

The variance as

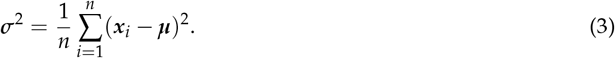

And the normalized output as

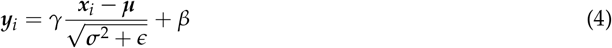

for *i* ∈ {1, …, *n*}. Then, it returns the output ***y*** = [***y***_1_, …, ***y***_*n*_]. The GELU activation is defined as:

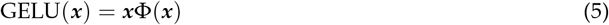

where Φ(***x***) is the cumulative distribution function of the standard normal distribution. To incorporate positional information, a learnable positional embedding is added to the input representation after the stem. This positional embedding ***P*** ∈ ℝ^1*×L×D*^ is initialized using a truncated normal distribution:

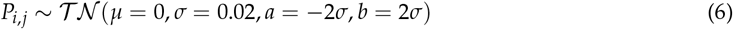

where 𝒯 𝒩 represents the truncated normal distribution with mean *µ* = 0, standard deviation *σ* = 0.02, and truncation bounds *a* = −2*σ* and *b* = 2*σ*. The output of the stem is then transposed to treat each time step as a separate “token,” enabling the subsequent layers to process the sequence effectively. The final output is a tensor ***X***_final_ ∈ ℝ^*B×L×D*^, where each of the *L* time steps in the sequence is represented as a “token” with *D* features.

#### Backbone

The backbone of Melchior processes the embedded input ***X***_final_ ∈ ℝ^*B×L×D*^, where *B* is the batch size, *L* = 4096 is the sequence length, and *D* = 768 is the embedding dimension. The backbone consists of *N* = 20 alternating Mamba and Transformer blocks:

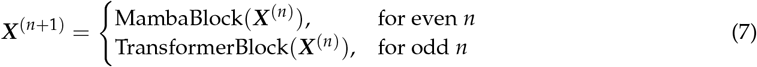

where *n* ∈ {0, 1, …, 18, 19} is the block index. Every such block follows a conventional residual formulation:

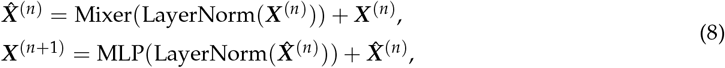

where a token mixing block is applied to the layer-normalized representation. Mamba blocks apply the MambaVision Mixer, while Transformer blocks employ multi-head self-attention. The MLP is a two-layer feed-forward network:

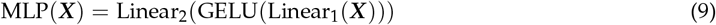

where Linear_1_ : ℝ^*D*^ → ℝ^*D*^_mlp_ and Linear_2_ : ℝ^*D*^_mlp_ → ℝ^*D*^, with *D*_mlp_ = 3072.

#### Head

The head processes the output of the backbone ***X***_*N*_ ∈ ℝ^*B×L×D*^ by applying a 1D adaptive average pooling to truncate to an output sequence length of *L*_out_ = 420, and then a linear projection to obtain the *C* = 5 class scores for each possible base. The final output is obtained by applying a log-softmax.

### 2.3 Micro Architecture

A state-space model (SSM) is defined as a linear-time invariant (LTI) system that projects an input stimulation *x*(*t*) ∈ ℝ to an output response *y*(*t*) ∈ ℝ through a hidden state *h*(*t*) ∈ 𝒞^*N*^. This system uses ***A*** ∈ ℝ^*N×N*^ as the evolution parameter and ***B, C*** ∈ ℝ^1*×N*^ as projection parameters. For continuous inputs, the system can be formulated as a group of linear ordinary differential equations:

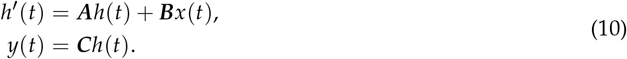

These parameters need to be discretized for further computational efficiency. Assuming a timescale Δ, Mamba uses the zero-order hold (ZOH) rule to obtain discrete parameters 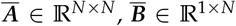, and 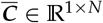:

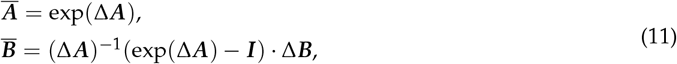

Then, (10) can be rewritten with the discrete parameters as:

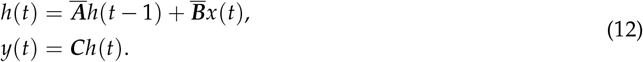

Lastly, a global convolution with kernel 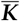 can be applied to compute (12):

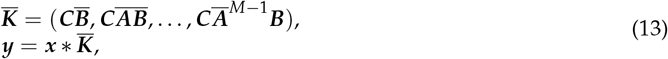

where *M* is the length of the input sequence ***x***.

#### 2.3.1 MambaVision Mixer

The token mixer proposed in [24] redesigns the original Mamba mixer by supplanting the causal convolution with the regular convolution, since causal convolutions limit the influence to one direction, which is unnecessary and restrictive for vision tasks. As shown in Figure 2, a symmetric branch without SSM was added, consisting of an additional convolution and SiLU activation, to compensate for any content lost due to the sequential constraints of SSMs. The embedded representations of both branches are then concatenated and projected by a final linear layer. Given an input ***X***_in_, MambaVision mixer calculates the output as:

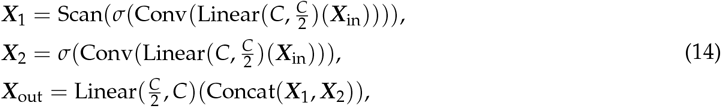

where Linear(*C*_in_, *C*_out_)(·) is a linear layer with *C*_in_ and *C*_out_ and input and output embedding dimensions, respectively, Scan(·) is the selective scan operation, Conv(·) is a 1D convolutional layer with a kernel size of 4, and *σ* is the SiLU [27] activation function.

**Figure 2:**
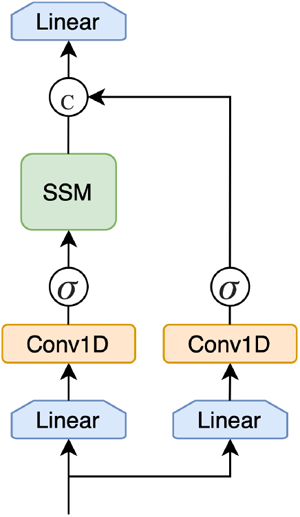
MambaVision Mixer uses convolutions and an SSM token mixer to enhance global context modeling. Notice that causal convolutions are replaced with their vanilla counterparts in the symmetric path.

#### 2.3.2 Multi-Head Self-Attention

The multi-head attention (MHA) operation is defined as:

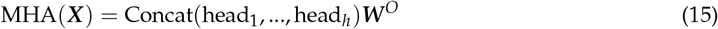

where each head is computed as:

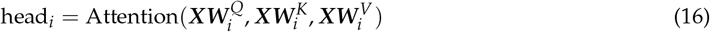

with 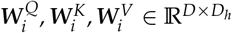 and ***W*** ^*O*^ ∈ ℝ^*D×D*^. Since we adopt *h* = 8 heads and the embedding dimension remains constant at *D*_model_ = 768, the head dimension is 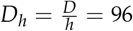. Attention is computed, without loss of generality, as:

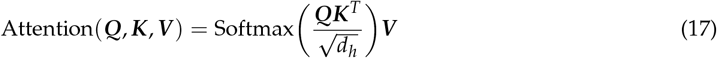

for queries, keys, and values ***Q, K***, and ***V***.

### 2.4 Data

For consistency of benchmarking, we adopt the dataset of reads collated by [13]. The training dataset comprises 116,072 reads from five different species: Arabidopsis thaliana, Homo sapiens [28], Caenorhabditis elegans [29], Escherichia coli [30], and synthetic constructs from Epinano [31]. Oxford Nanopore Technology’s R9.4 and R9.4.1 flow cells were used to generate the dataset. The Arabidopsis data was derived by [13] from 17-day-old seedlings, while other species’ data were obtained from publicly available sources. The training dataset consists of 24,370 Arabidopsis reads aligned to the Araport11 transcriptome, 29,728 Epinano synthetic construct reads aligned to their released reference, 30,048 Homo sapiens reads aligned to the gencode v33 transcriptome, 24,192 Caenorhabditis elegans reads aligned to the CE11 transcriptome, and 7,734 Escherichia coli reads aligned to the transcriptome generated from the NCBI assembly database. For training and validation purposes, reads were randomly selected and segmented into chunks of 4096 signal values. The raw input signals were normalized by median absolute deviation.

The test dataset consists of 100,000 reads from five different species, including Homo sapiens [28], Arabidopsis thaliana [32], Mus musculus [33], S. cerevisiae [34], and Populus trichocarpa [35]. These reads were basecalled with Guppy v4.4.0 and aligned to their respective transcriptomes. Any reads that aligned to the mitochondrial genome were discarded.

### 2.5 Training

Melchior was trained on an NVIDIA A100-SXM4 GPU with 80 GB of RAM for 223 hours, completing 16 epochs. We used CTC loss with label smoothing and managed the learning rate through an initial 5% linear warm-up to 6e-4 followed by cosine decay. Training utilized a batch size of 32 with stochastic dropout. Stochastic weight averaging was applied to improve model generalization and stability. The final model was 134 million parameters. During inference, a batch size of 128 was adopted for all models.

### 3 Results & Discussion

We compare Melchior to other open ONT basecallers on the basis of percent identity and error profiles. Our analysis reveals that Melchior outperforms existing state-of-the-art basecallers across multiple species. Table 1 presents a comparison of the performance metrics between Guppy, RODAN, GCRTCall, and Melchior for the five species. Melchior achieves the highest identity for all species, with notable improvements in human (94.40%) and Arabidopsis (94.13%) samples. Remarkably, Melchior demonstrates the lowest error rates in nearly all categories. It excels in minimizing insertion rates, with the best performance across all species, particularly in mouse samples (1.74%). Melchior also leads in reducing mismatch rates, showing significant improvements over other basecallers, especially in human (1.17%), mouse (2.78%), and Arabidopsis (1.27%) datasets.

**Table 1:** Performance comparison between Guppy, RODAN, GCRTCall, and Melchior.

| Species | Basecaller | Identity (%) | Insertion rate (%) | Deletion rate (%) | Mismatch rate (%) |
| --- | --- | --- | --- | --- | --- |
| Yeast | Guppy | 91.41 | 2.31 | 3.77 | 2.36 |
|  | RODAN | 91.41 | 2.08 | 3.69 | 2.55 |
|  | GCRTcall | 92.52 | 2.01 | **3.08** | 2.26 |
|  | Melchior | **92.60** | **2.00** | 3.11 | **2.09** |
| Human | Guppy | 90.64 | 1.99 | 4.88 | 2.27 |
|  | RODAN | 93.19 | 1.54 | 3.16 | 1.53 |
|  | GCRTcall | 94.13 | 1.48 | 2.72 | 1.37 |
|  | Melchior | **94.40** | **1.45** | **2.53** | **1.17** |
| Mouse | Guppy | 87.74 | 2.51 | 5.90 | 3.63 |
|  | RODAN | 88.02 | 1.94 | 6.14 | 3.61 |
|  | GCRTcall | 89.63 | 1.88 | 5.10 | 3.17 |
|  | Melchior | **90.17** | **1.74** | **5.04** | **2.78** |
| Poplar | Guppy | 90.22 | 2.50 | 4.43 | 2.61 |
|  | RODAN | 91.11 | 2.26 | 3.82 | 2.47 |
|  | GCRTcall | 92.14 | 2.10 | 3.31 | 2.22 |
|  | Melchior | **92.42** | **2.07** | **3.28** | **1.95** |
| Arabidopsis | Guppy | 91.61 | 1.92 | 4.30 | 2.01 |
|  | RODAN | 92.90 | 1.59 | 3.41 | 1.69 |
|  | GCRTcall | 93.91 | 1.52 | 2.84 | 1.50 |
|  | Melchior | **94.13** | **1.49** | **2.77** | **1.27** |

We also evaluated the basecallers’ performance in terms of read alignment rates, which provide insight into the overall usability of the called sequences. Melchior demonstrates superior performance with a mean of 99,253 aligned reads, surpassing GCRTCall (98,763), Guppy (98,107), and RODAN (98,034). This represents a 0.5% improvement over the next best performer, GCRTCall, and a notable 1.2% increase compared to Guppy. Figure 3 presents the mean number of aligned reads across all species for each basecaller.

**Table 2.**
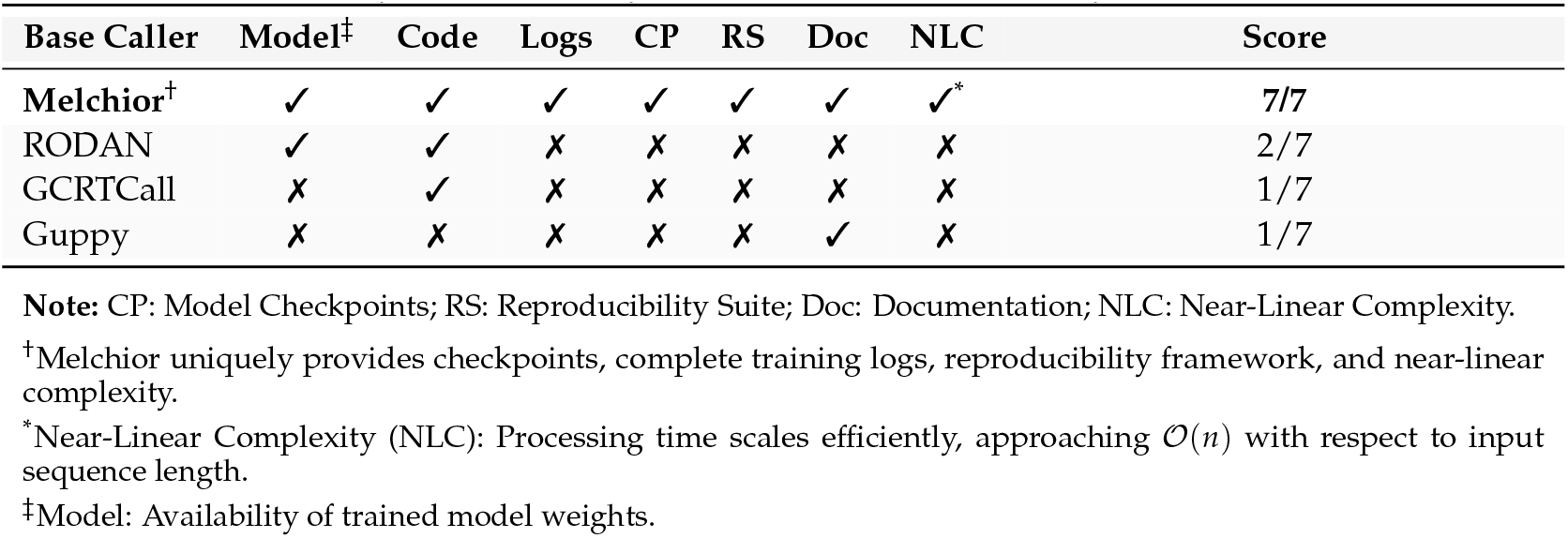
Transparency, Reproducibility, and Computational Efficiency of Nanopore Base Callers.

**Figure 3:**
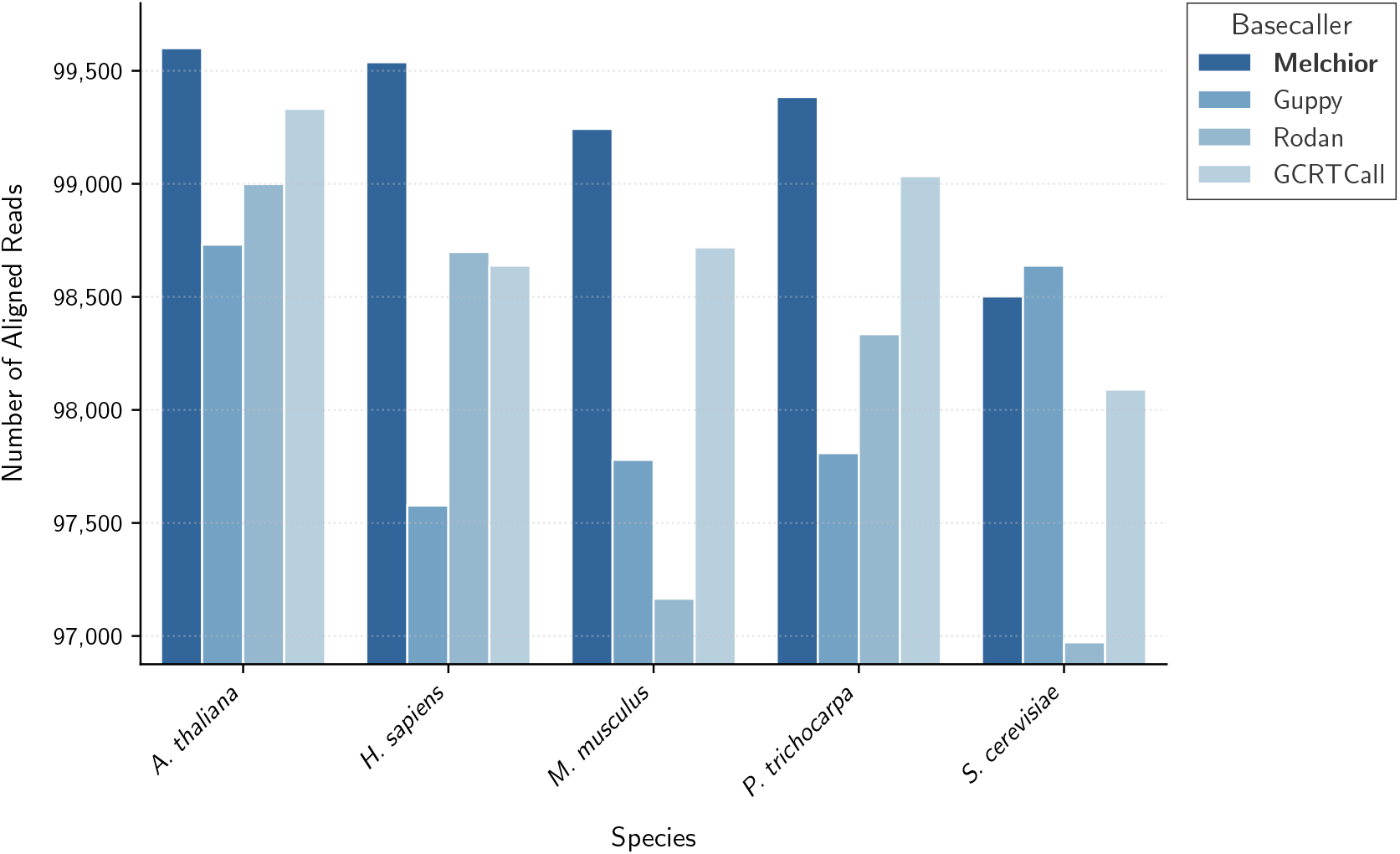
Comparison of aligned reads across species and basecallers. Melchior demonstrates consistently superior alignment rates across all tested species, with particular improvement in challenging datasets.

Moreover, as shown in Figure 4, Melchior excels in the alignment length distribution, with a higher percentage of reads having an alignment length greater than or equal to 1000 base pairs (bp). The alignment length distribution directly impacts the accuracy and completeness of genome assemblies, the detection of structural variations, and the resolution of complex genomic regions; longer alignments typically indicate higher-quality reads and more reliable mapping.

**Figure 4:**
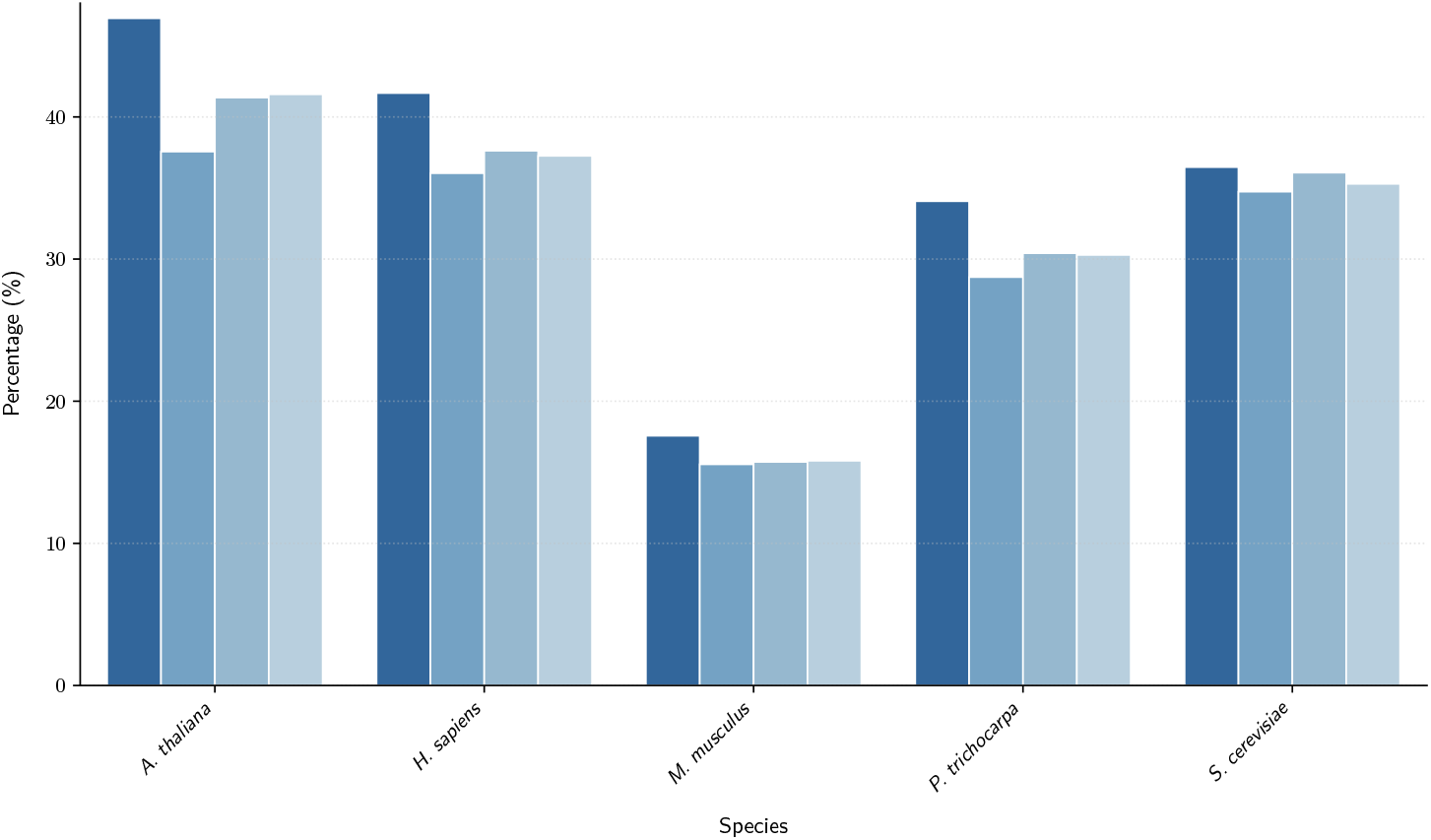
Alignment Length Distribution (≥1000bp) for different basecallers and species. Refer to legend from Figure 3.

Beyond the generally advantageous reads produced by Melchior, its exceptional strengths are particularly evident in homopolymer calling. Combining single-base-pair resolution with long-range contextual information, Melchior is able to correctly identify and resolve regions of high repetition and low sequence complexity. Figure 5 depicts a series of heatmaps, where each heatmap shows the percentage of high-quality reads for each basecaller and species at a given minimum homopolymer length. We defined high-quality reads as those with a mapping quality score of 30 or higher. Melchior consistently outperforms other basecallers across all minimum homopolymer lengths (50, 75, 100, and 125), surpassing Guppy and RODAN by tens of orders of magnitude and outperforming GCRTCall by a factor of several fold.

**Figure 5:**
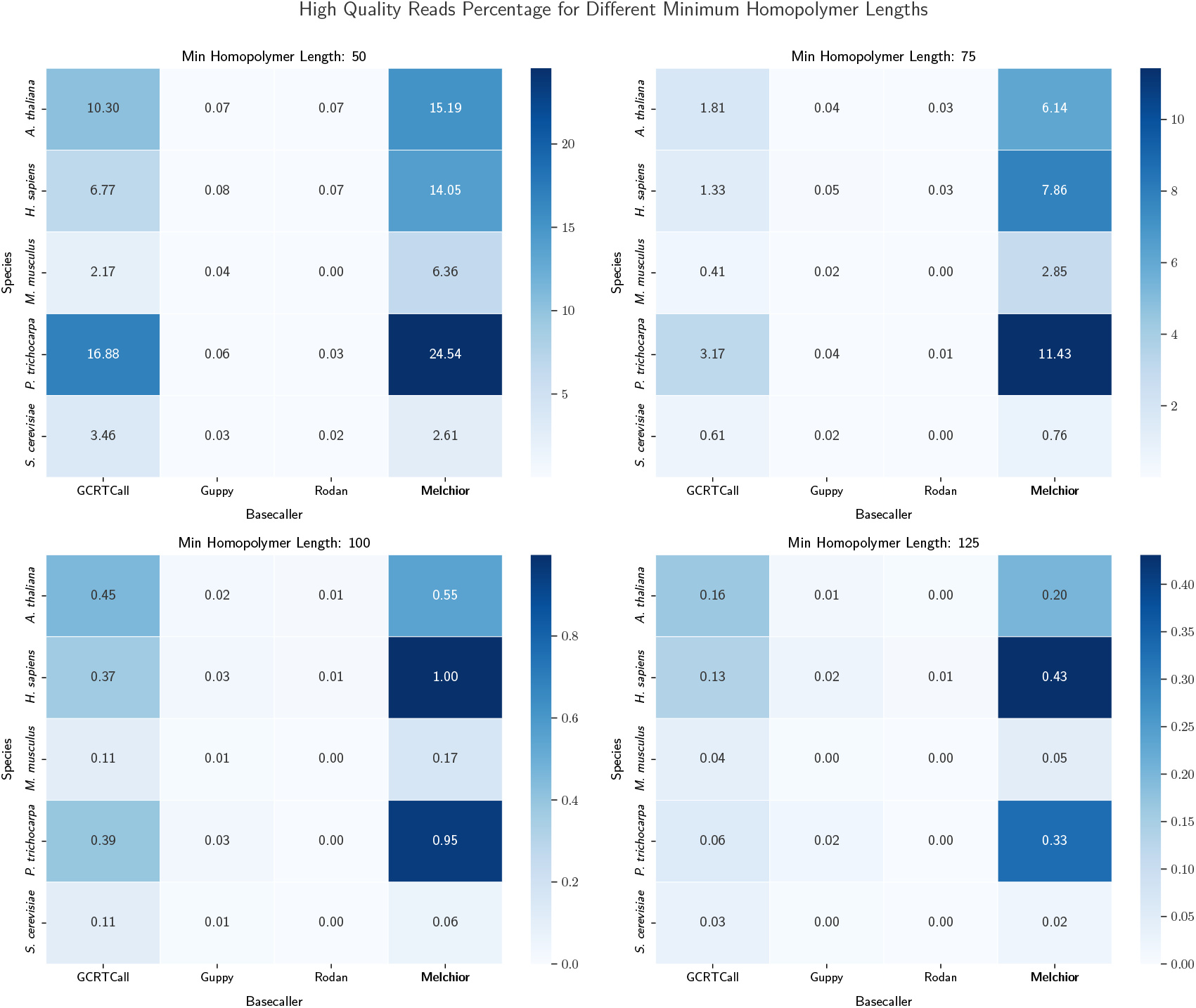
Comparison of high-quality read percentages for different basecallers and species, stratified by minimum homopolymer length. Melchior consistently outperforms other basecallers, with marked improvements in high-quality read recovery for large homopolymers and challenging datasets.

In addition to achieving state-of-the-art basecalling performance, Melchior sets a new standard with respect to openness and efficiency. The model is fully reproducible, with all training details, logs, evaluation scripts, and checkpoints made public. Melchior also achieves nearly linear complexity by virtue of its hybrid backbone. This represents a significantly more efficient sequencing paradigm than traditional quadratic-time basecallers. Table 2 illustrates these differences.

## 4 Conclusion

We introduced Melchior, a hybrid Mamba-Transformer RNA basecaller. Our analysis demonstrates that a hybrid architecture achieves the best of both worlds: high-accuracy basecalling and efficiency, outperforming existing state-of-the-art methods. Melchior was designed with reproducibility and accessibility in mind; we hope that its efficiency and ease of use will help unlock new applications in field sequencing, democratize the generation of large-scale sequencing data, and become a valuable resource for the community.

Furthermore, we demonstrated Melchior’s ability to handle local sequence challenges while maintaining its strength in global read characteristics. As such, this paper represents the first principled approach to homopolymer calling, without any redundancies involving depth of coverage. Accurate homopolymer calling is crucial for precise genome assembly and variant detection, especially in repetitive regions or areas with microsatellite instability or poly-A tails. We believe these insights will inspire new efforts to hybridize architectures in sequencing pipelines.

## 5 Code Availability Statement

The complete implementation of Melchior is publicly available at github.com/elonlit/melchior. The repository includes training scripts, model architectures, and evaluation code used in this work. Full training logs, experimental metrics, and hyperparameter configurations are hosted at wandb.ai/julian-q/melchior for reproducibility purposes. Trained model weights and checkpoints can be found at hugging-face.com/elonlit/Melchior.

## Acknowledgements

We thank Julian Quevedo for insightful discussions and providing computational resources, which greatly facilitated the development and testing of Melchior.

